# Specific detection of cell-free DNA derived from intestinal epithelial cells using methylation patterns

**DOI:** 10.1101/409219

**Authors:** Roni Lehmann-Werman, Aviad Zick, Cloud Paweletz, Marisa Welch, Ayala Hubert, Miriam Maoz, Tal Davidy, Judith Magenheim, Sheina Piyanzin, Daniel Neiman, Joshua Moss, Hadar Golan, Eran Israeli, Matan Fischer, Eran Segal, Markus Grompe, Alon Pikarsky, Talia Golan, Benjamin Glaser, Ruth Shemer, Brian Wolpin, Yuval Dor

## Abstract

Epithelial cells of the intestine undergo rapid turnover and are thought to be cleared via stool. Disruption of tissue architecture, as occurs in colorectal cancer (CRC), results in the release of material from dying intestinal epithelial cells to blood. This phenomenon could be utilized for diagnosis and monitoring of intestinal diseases, if circulating cell-free DNA (cfDNA) derived from intestinal cells could be specifically identified. Here we describe two genomic loci that are unmethylated specifically in intestinal epithelial cells, allowing for sensitive and specific detection of DNA derived from such cells. As expected, intestinal DNA is found in stool, but not in plasma, of healthy individuals. Patients with inflammatory bowel disease (IBD) have minimal amounts of intestinal cfDNA in the plasma, whereas patients with advanced CRC show a strong signal. The intestinal markers are not elevated in plasma samples from patients with pancreatic ductal adenocarcinoma (PDAC), and a combination of intestine- and pancreas-specific markers allowed for robust differentiation between plasma cfDNA derived from CRC and PDAC patients. Intestinal DNA markers provide a mutation-independent tool for monitoring intestinal dynamics in health and disease.

## Introduction

Reliable biomarkers for detection and monitoring of intestinal diseases are missing in the toolbox of gastroenterologists and oncologists. While colonoscopy and fecal screens are useful for early detection of CRC, compliance to these tests remains suboptimal. Circulating biomarkers for CRC and IBD exist and are in clinical use, for example carcinoembryonic antigen (CEA), methylated SEPT9 and c-reactive protein, but their sensitivity and specificity are limited (Nikolaou et al., 2018; Sands, 2015). Beyond pathology, our ability to study intestinal dynamics in healthy people, during development and in different physiological conditions, is minimal.

Cell-free DNA (cfDNA) released from dying cells is emerging as an important substrate for liquid biopsies, used for detection and monitoring of various clinical situations, including non-invasive prenatal testing (NIPT)(Bianchi and Chiu, 2018; Wong and Lo, 2016), identification of somatic mutations in the plasma of cancer patients (Wan et al., 2017), and early detection of graft rejection in organ transplant recipients (Snyder et al., 2011). All of these are based on genetic differences between normal tissue and the tissue of interest (Aravanis et al., 2017; Norwitz and Levy, 2013; Snyder et al., 2011; Wan et al., 2017). While the detection of somatic mutations in cancer in general, and CRC in particular, is revolutionizing diagnostic oncology, there are important limitations to this approach. For example, individual colon cancers have different mutations, which are dynamic and often sub-clonal, and hence require a large panel of markers to be tested. In the context of early cancer detection, the identification of an oncogenic mutation in cfDNA does not reveal the location of cancer (Aravanis et al., 2017). More fundamentally, assays based on somatic mutations are blind to cfDNA derived from tissues having a normal genome, as occurs in healthy individuals, in patients with non-malignant pathologies, and in bystander tissue damage in cancer.

We and others have recently developed an approach for detecting the tissue origins of cfDNA, based on tissue-specific methylation patterns which are typical to each cell type, are conserved among cells of the same type across individuals, and largely maintained with age as well as in pathological conditions (Dor and Cedar, 2018; Guo et al., 2017; Lehmann-Werman et al., 2016; Sun et al., 2015). We performed comparative methylome analysis to identify genomic loci with a methylation pattern that is typical to a specific cell type, and detected these patterns in the plasma. Using bisulfite treatment of cfDNA followed by PCR and sequencing or alternatively digital droplet PCR, we demonstrated sensitive and specific detection of cfDNA derived from the exocrine and endocrine pancreas, liver, heart and brain (Gala-Lopez et al., 2018; Lehmann-Werman et al., 2018; Lehmann-Werman et al., 2016; Zemmour et al., 2018).

Here we characterize two methylation markers of intestinal epithelial cells, document their sensitivity and specificity in detection of intestinal DNA, and report the levels of DNA derived from intestinal epithelial cells in the plasma and stool of healthy individuals as well as in the context of intestinal pathologies.

## Results

### Identification of intestine-specific methylation markers

To identify genomic loci that show an intestine-specific methylation pattern, we compared the methylome of human colon DNA to a panel 20 human tissue and cell type methylomes obtained from public sources or generated in our lab (Moss et al, manuscript in review) using the Illumina Infinium HumanMethylation450 BeadChip array. We focused on CpG sites that were differentially methylated, namely unmethylated in colon DNA and methylated elsewhere. Multiple sites fulfilled our criteria of having a methylation index of <0.5 in colon DNA and an index of >0.8 in the blood (Figure 1a). We selected for further analysis two of the top hits, one located in the 3′ UTR region of the ECH1 gene and one located in an unspecified locus that we named Col (Supplemental Figure S1) (see genomic coordinates in Methods). To take advantage of the information contained in methylation haplotype blocks (Guo et al., 2017; Lehmann-Werman et al., 2016) we defined “expanded windows” around the marker cytosine in each locus, containing additional CpG sites; the ECH amplicon was 115 bp long and contained 5 CpG sites; the Col amplicon was 118 bp long and contained 8 CpG sites. We treated genomic DNA from multiple tissues with bisulfite, PCR amplified the two expanded windows, and sequenced the products on a NextSeq platform to determine the tissue-specific methylation status of all cytosines in each amplicon. As shown in Figure 1B, for both markers we identified fully unmethylated molecules (that is, unmethylated in all 5 or 8 CpG sites in the ECH1 or Col amplicon respectively) only in intestinal DNA. Strikingly, the DNA from sorted colon epithelial cells was almost completely demethylated in these loci, indicating that these are true specific markers of the intestinal epithelium and not another compartment in intestinal tissue (Figure 1B).

**Figure 1:**
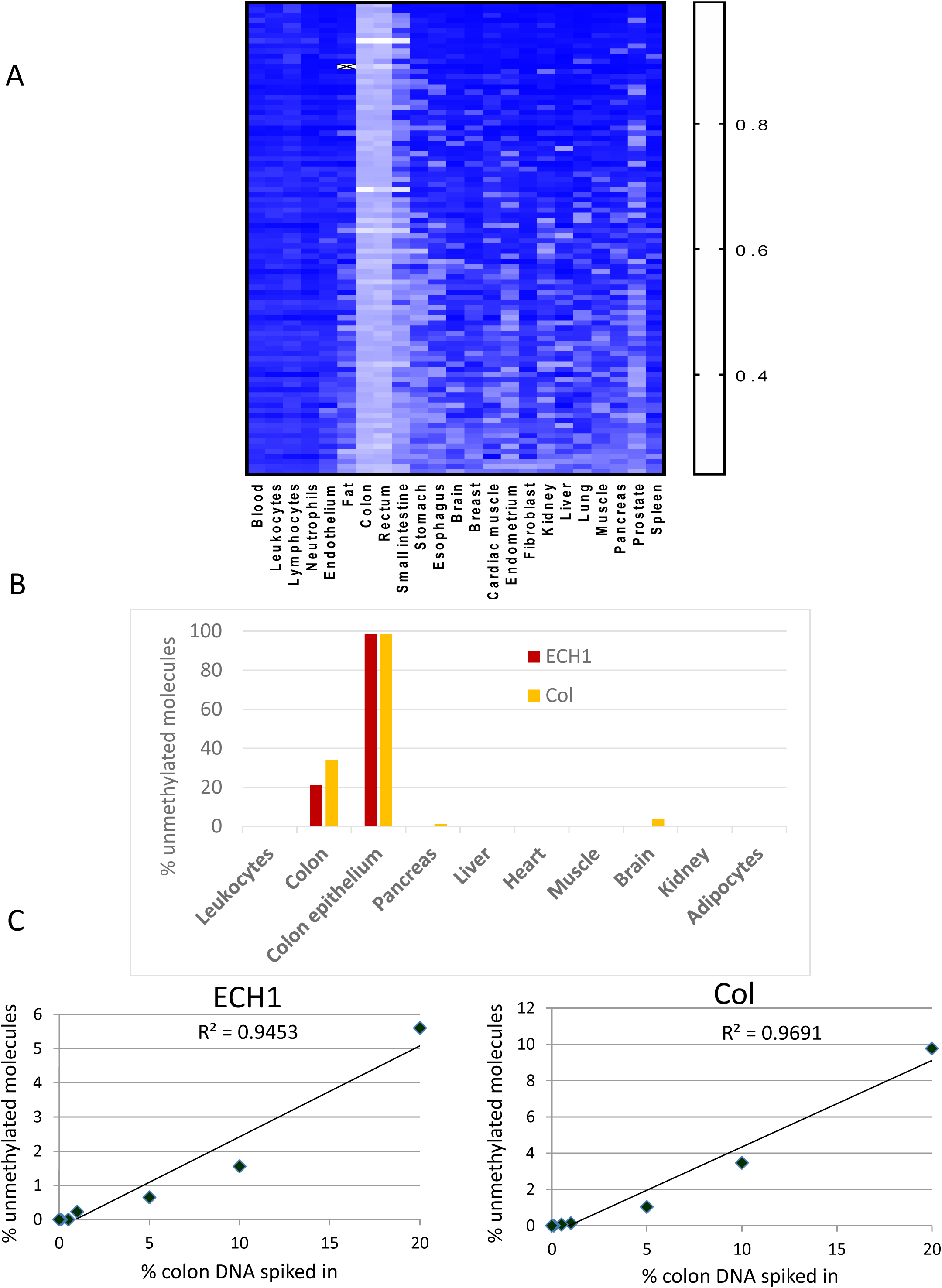
Intestine-specific markers. **A.** Comparative tissue methylome analysis reveals multiple cytosines that are methylated or unmethylated specifically in the intestine. Shown is a heatmap of the top 100 cytosines distinguishing intestinal DNA from all other tissues. Y axis represents the 90^th^ percentile of methylation for the colon and the 10^th^ percentile of methylation fraction for all other tissues. **B.** Assay specificity. Methylation status of haplotype blocks near the ECH1 and Col loci in DNA from multiple tissues. Shown is the percentage of molecules in which all CpG sites were unmethylated. **C.** Assay sensitivity and accuracy in vitro. DNA from healthy human colon was mixed with blood DNA as indicated, and the fraction of molecules unmethylated in the ECH2 or Col markers was determined.

To assess sensitivity and accuracy of intestinal DNA detection, we diluted colon DNA into whole blood DNA, and used the methylation assay to assess the relative contribution of colon DNA to the mix. Analysis of both markers allowed accurate estimation of the fraction of colon DNA in the mixture (Figure 1C). We were able to distinguish as little as 0.1% colon DNA, or 23 colon genomes mixed with 23,000 blood genomes (Figure 1C).

These findings establish two genomic loci, each containing a block of methylation sites, which are unmethylated specifically in intestinal epithelial cells and can be detected in a sensitive manner in DNA mixtures.

### Intestinal cfDNA in healthy individuals

We measured intestine-derived cfDNA in the plasma of healthy individuals. As expected, healthy individuals showed a minimal signal, consistent with removal of dying intestinal cells to the lumen of gut rather than to blood (Figure 2A). To test this idea directly, we examined DNA extracted from stool samples of healthy individuals. The vast majority of stool DNA is derived from bacteria, as observed in next generation sequencing (data not shown). In most samples, no human DNA could be detected. However, when we examined a subset of stool DNA samples that did contain human DNA (n=10), we found that a large fraction of the human DNA molecules in most samples (7/10) carried the intestinal pattern of methylation (Figure 2B). Thus, intestinal DNA markers in healthy plasma and stool reflect the established route of clearance of intestinal DNA, via the lumen of the gut.

**Figure 2:**
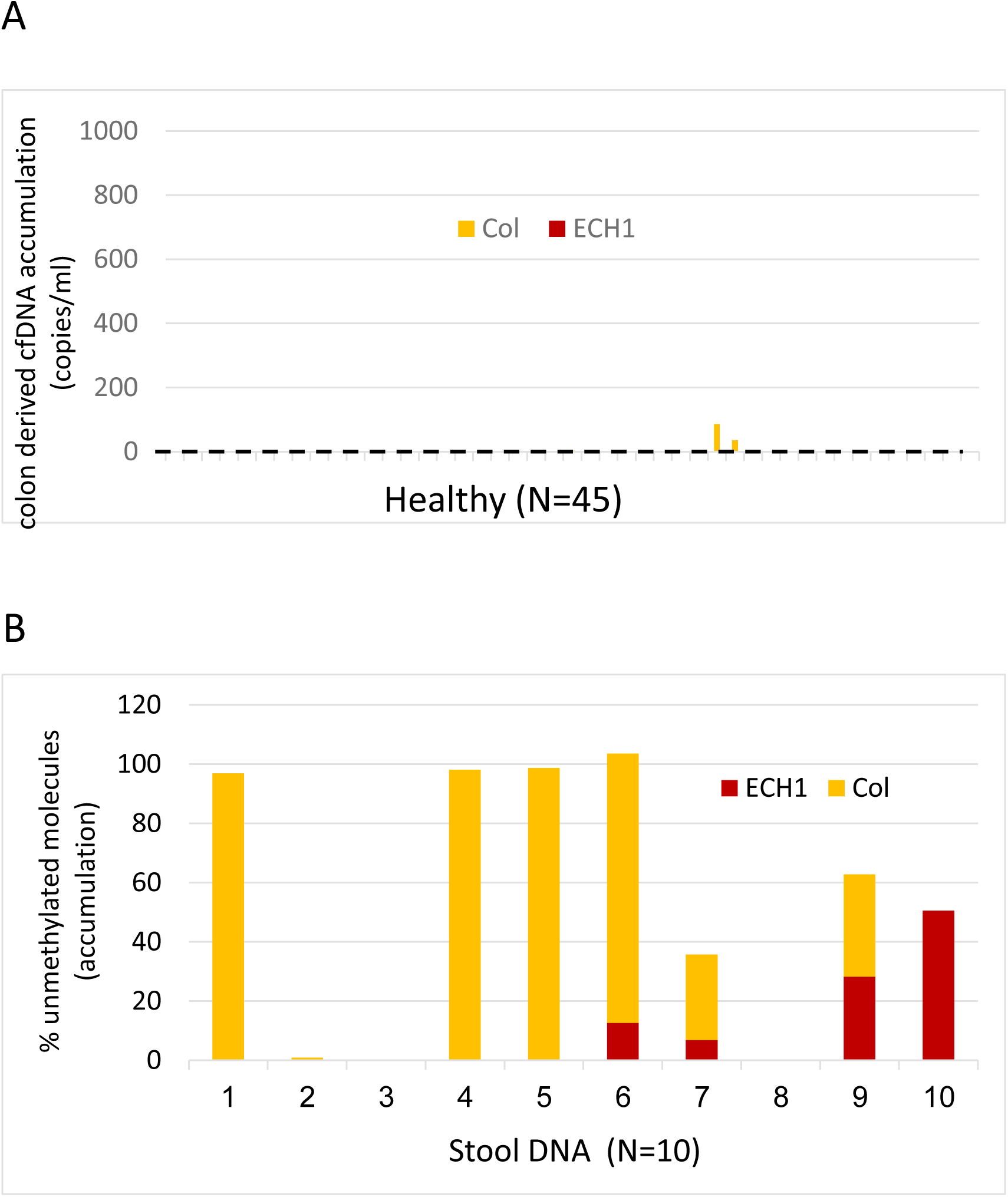
Intestine-derived DNA in healthy individuals. **A.** Concentration of intestinal cfDNA in the plasma of 45 healthy donors. The concentration was measured by multiplying the fraction of colon cfDNA by the concentration of total cfDNA. Dashed line indicates median (95% CI: 0,0). **B.** Intestinal DNA in stool samples. Shown are the results of analysis from 10 stool DNA samples from healthy donors that were used in a microbiome analysis, and found to contain 0.04-0.2% human DNA.

### Intestinal cfDNA in patients with IBD or CRC

We assessed the level of intestinal cfDNA in the plasma of patients with inflammatory bowel disease during remission (n=16; see patient details, Supplementary Table S1). Surprisingly, IBD patients had a minimal baseline intestinal cfDNA signal that was indistinguishable from that of healthy individuals (Figure 3A). In stark contrast, most patients with advanced CRC (42 out of 50, 84%) had a strong intestinal signal (Figure 3C). Plasma from CRC patients with a local disease (n=20) did not show a signal above baseline (Figure 3B). The area under the curve of the ROC curve for all CRC patients compared to healthy controls and IBD patients was 0.84 (Figure 3D).

**Figure 3:**
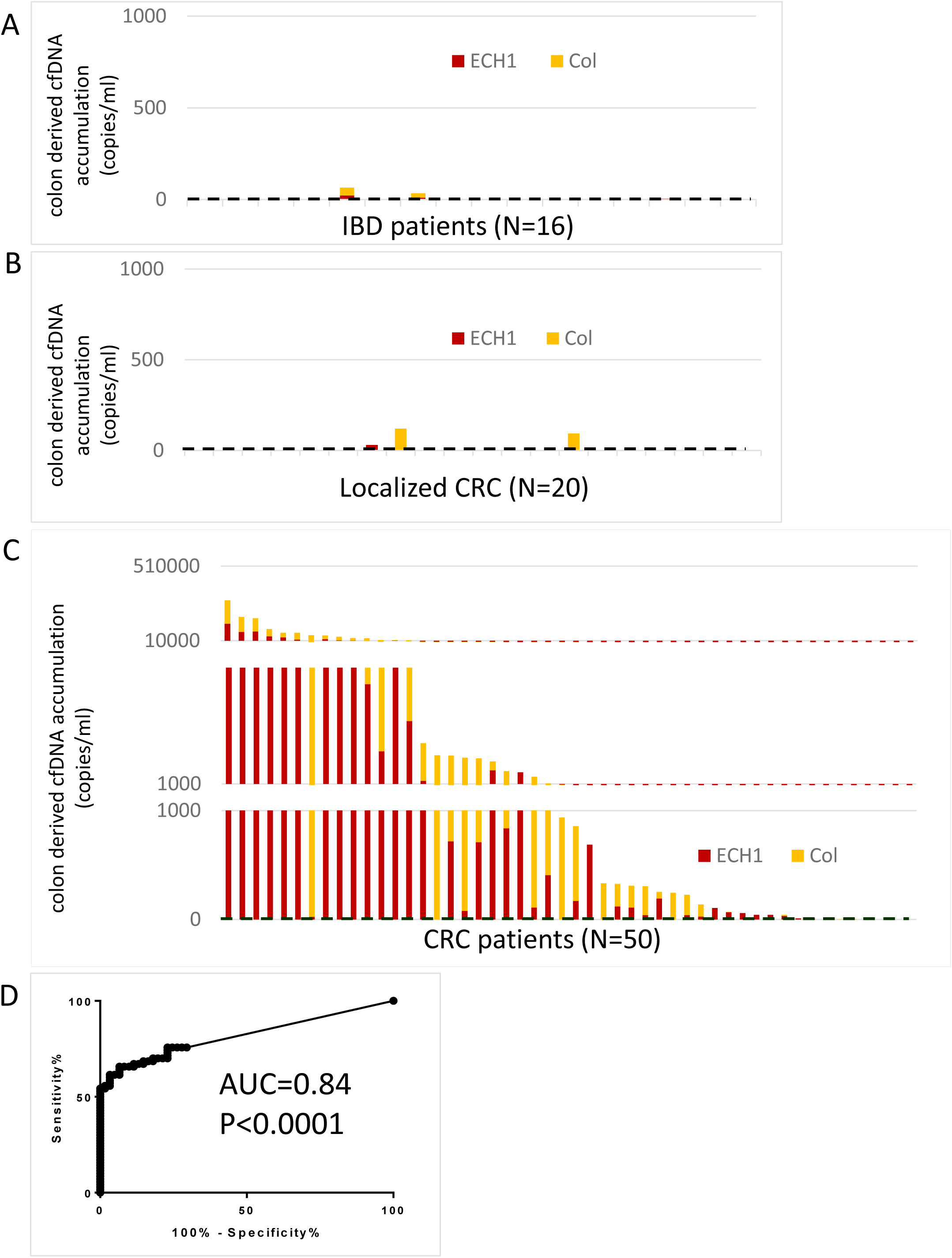
Intestinal cfDNA in IBD and CRC patients. **A.** Concentration of intestinal cfDNA in the plasma of 16 IBD patients in various stages of disease. The concentration was measured by multiplying the fraction of intestinal cfDNA by the concentration of total cfDNA. **B.** Intestinal cfDNA in the plasma of 20 localized CRC patients. The concentration was measured by multiplying the fraction of intestinal cfDNA by the concentration of total cfDNA. **C.** Intestinal cfDNA in the plasma of 50 advanced CRC patients. The concentration was measured by multiplying the fraction of intestinal cfDNA by the concentration of total cfDNA. **D.** ROC curve of all CRC samples (localized and metastatic) vs. all non-CRC samples (healthy and IBD).

Thus, intestinal DNA reaches the blood in advanced cancer, but not in healthy conditions or in remitting inflammatory bowel disease, suggesting that the methylation assay can distinguish between intestinal damage due to cancer or to other conditions. We note that 2/16 IBD samples showed a signal compared with 2/45 healthy samples (Figure 2A, 3A), suggesting that a more sensitive assay on a larger cohort may reveal an intestinal cfDNA signal in IBD (see discussion).

### Distinguishing plasma of patients with metastatic colorectal or metastatic pancreatic cancer

To test assay specificity *in vivo*, we assessed a blinded collection of plasma samples from patients with either metastatic CRC (N=38) or metastatic PDAC (N=12). We designed new specific methylation markers for the exocrine pancreas (that is, unmethylated specifically in the exocrine pancreas and methylated elsewhere; Supplemental figure S2), similar to those previously published (Lehmann-Werman et al., 2016) but ensuring that the markers are fully methylated particularly in intestinal epithelium. We established a multiplex PCR protocol that allowed for co-amplification of 2 intestinal markers and 2 pancreatic markers from the same cfDNA sample, and tested all markers on the blinded samples. As shown in Figure 4, the intestine and pancreas markers showed an excellent differentiation between plasma samples of the two groups of patients, supporting utility of the approach for rapid identification of the tissue of origin of circulating DNA.

**Figure 4:**
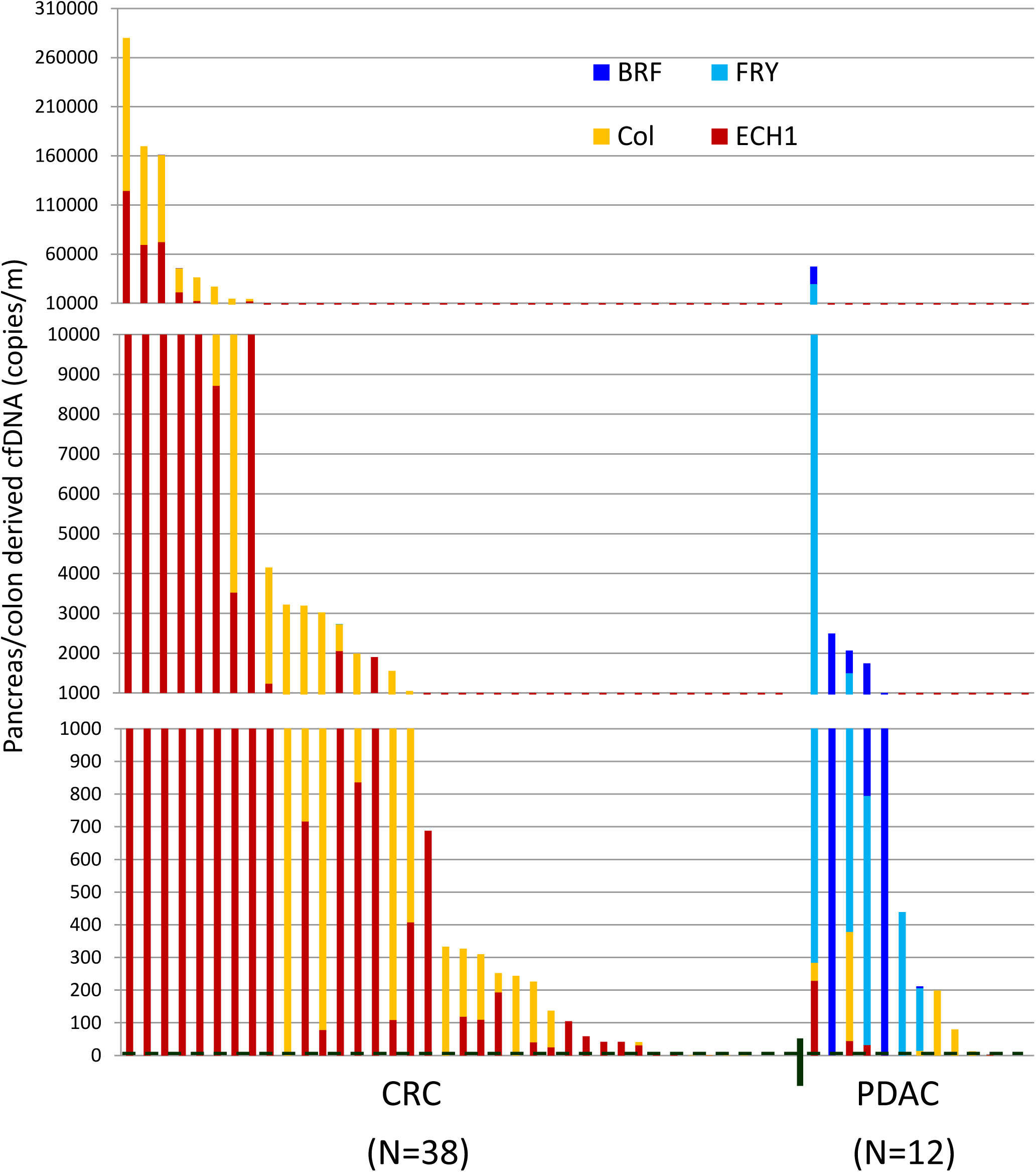
Differentiation between CRC and PDAC. Intestinal and exocrine pancreas cfDNA in the plasma of 38 CRC patients and 13 PDAC patients. Intestinal markers ECH1 and Col are shown in blue and light-blue respectively. Pancreas markers BRF and FRY are shown in green and light-green respectively. The concentration was measured by multiplying the fraction of intestinal cfDNA by the concentration of total cfDNA.

## Discussion

We have described two DNA methylation markers that can be used to identify intestinal DNA when mixed with DNA from other sources, based on bisulfite conversion, PCR amplification and massively parallel sequencing.

Our findings are consistent with the known route of clearance of dying intestinal cells under healthy conditions, to the lumen of the gut and then to stool. We hypothesize that the stool contains mostly DNA from the distal gut, and that DNA from the small intestine is degraded before exiting. Identification of methylation markers distinguishing between the small and large intestine will allow more specific tracing of the origins of human DNA in stool.

Surprisingly, we found very little intestinal DNA in the plasma of remitting IBD patients. More work is needed to determine if intestinal DNA reaches the circulation in relapsing IBD, as would be expected given the widespread tissue damage in this condition. In contrast, intestinal DNA was readily detected in the plasma of patients with advanced disseminated (but not localized) CRC. Moreover, combining the intestinal markers with two novel exocrine pancreas methylation markers accurately distinguished between plasma from CRC and PDAC patients.

These findings provide a proof of concept that methylation patterns can be used to detect cfDNA derived from intestinal epithelial cells. Two recent studies have reached similar conclusions, using different markers and different detection techniques. Guo et al used deconvolution of the plasma methylome to identify patients with advanced CRC (Guo et al., 2017), and Gai et al used digital droplet PCR to identify colon DNA in the plasma of CRC patients (Gai et al., 2018). While a direct comparison between the methods was not performed, our directed PCR-sequencing assay, which captures methylation haplotype blocks in specific loci in great depth, is likely more sensitive and specific than the other methods.

We have identified DNA methylation markers that are highly specific to DNA originating from intestinal epithelial cells, with no evidence for a signal derived from other tissues. However, the limitation of our current assay sensitivity does not permit the detection of intestinal DNA in the plasma of patients with localized CRC, which will be essential for the development of early diagnosis approaches. We propose that by testing multiple independent intestinal markers on the same cfDNA sample, assay sensitivity can be dramatically enhanced, potentially allowing the detection of disease at a milder, earlier stage. Our comparative methylome analysis revealed many dozens of genomic loci that could serve as intestinal markers. Alternatively, it is possible that the failure to detect intestinal cfDNA in localized CRC may reflect biology- the fact that epithelial cell turnover in early stages of the CRC (and perhaps also in remitting IBD) releases material to stool rather than to plasma. More generally, the route of clearance of DNA from dying cells in different tissues, in health and disease, might be an unappreciated determinant of cfDNA biology.

## Materials and Methods

### Patients

All clinical studies were approved by the ethics committees of the Hadassah Hebrew-University Medical Center, Sheba Medial Center and Dana Farber Cancer Institute.

### Biomarkers

Tissue-specific methylation biomarkers were selected after a comparison of publicly available genome-wide DNA methylation datasets generated using Illumina Infinium HumanMethylation450k BeadChip array. The ECH1 marker contains 5 CpG dinucleotides, included in a 115 bp fragment surrounding position CG12028674, amplified from bisulfute-treated DNA using primers GTTAGAAGGTATAGAAATAATTGTTAT and TCTCCAAACTCTAAAAACCCT.

The Col marker contains 8 CpG dinucleotides, included in a 118 bp fragment surrounding position CG15139063, amplified from bisulfute-treated DNA using primers GGTGGGTATGTGATTTGTGA and TAACCCCTAAATCCAAAACC.

### Sample Preparation and DNA Processing

Blood samples were collected in EDTA-containing plasma-preparation tubes and centrifuged for 10 min in 4 degree at 1,500× g. The supernatant was transferred to a fresh 15 ml conical tube without disturbing the cellular layer and centrifuged again for 10 min in 4 degrees at 3000 × g. The supernatant was collected and stored in −80c.

Cell-free DNA was extracted from 1-4 mL of plasma using the QIAsymphony liquid handling robot (Qiagen) and treated with bisulfite (Zymo Research). DNA concentration was measured using Qbit double-strand molecular probes (Invitrogen). Bisulfite-treated DNA was PCR amplified using primers specific for bisulfite-treated DNA but independent of methylation status at monitored CpG sites.

Primers were bar-coded, allowing the mixing of samples from different individuals when sequencing products. Sequencing was performed on PCR products using MiSeq Reagent Kit v2 (MiSeq, Illuminamethod) or NextSeq 500/550 v2 sequencing reagent kits. Sequenced reads were separated by barcode, aligned to the target sequence, and analyzed using custom scripts written and implemented in Matlab. Reads were quality filtered based on Illumina quality scores. Reads were identified by having at least 80% similarity to target sequences and containing all the expected CpGs in the sequence. CpGs were considered methylated if “CG” was read and were considered unmethylated if “TG” was read. The efficiency of bisulfite conversion was assessed by analyzing the methylation of non-CpG cytosines.

### Colon epithelium sorting

Fresh surgical samples of the colon were dissociated to single cells as described (Roche, 2001), immunostained using anti-CD45 eFluor 450, anti-CD31 eFluor 450, anti-CD234A eFluor 450 (eBioscience) and anti-CD326 (Miltenyi) antibodies, and sorted to isolate cells from which genomic DNA was prepared.

## Acknowledgements

Supported by grants from the National Cancer Institute (pancreatic cancer early detection consortium), Human Islet Research Network of the NIH (HIRN), The Ernest and Bonnie Beutler Research Program of Excellence in Genomic Medicine, the Juvenile Diabetes Research Foundation, the Kahn Foundation, and the Alex U. Soyka pancreatic cancer fund.

## Legend to Supplemental Figures

**Supplemental Figure S1:**
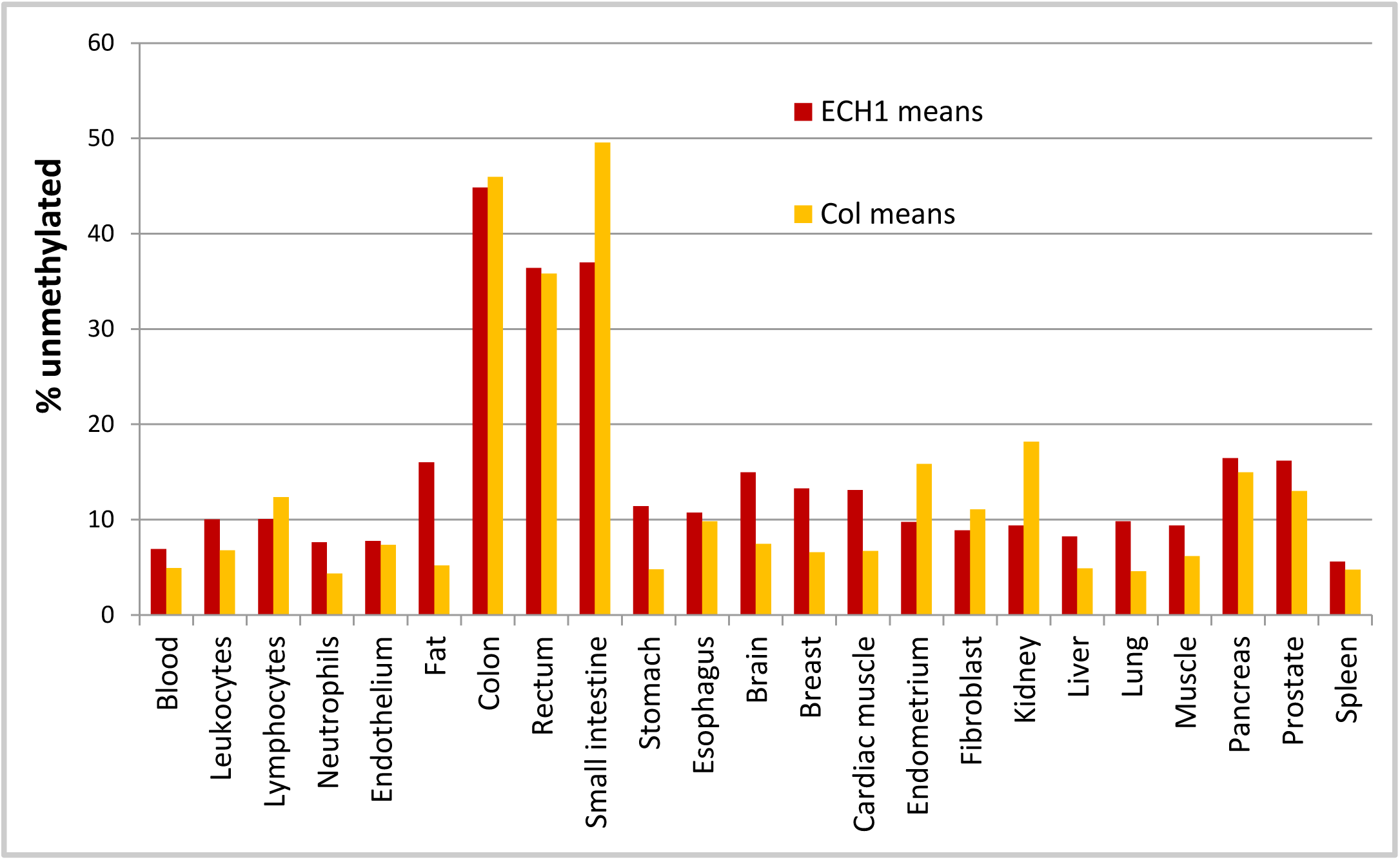
Identification of intestine-specific DNA methylation markers. Methylation status of the individual CpG sites at the ECH1 and Col loci that are captured in the Illumina 450k array, across multiple tissues.

**Supplemental Figure S2:**
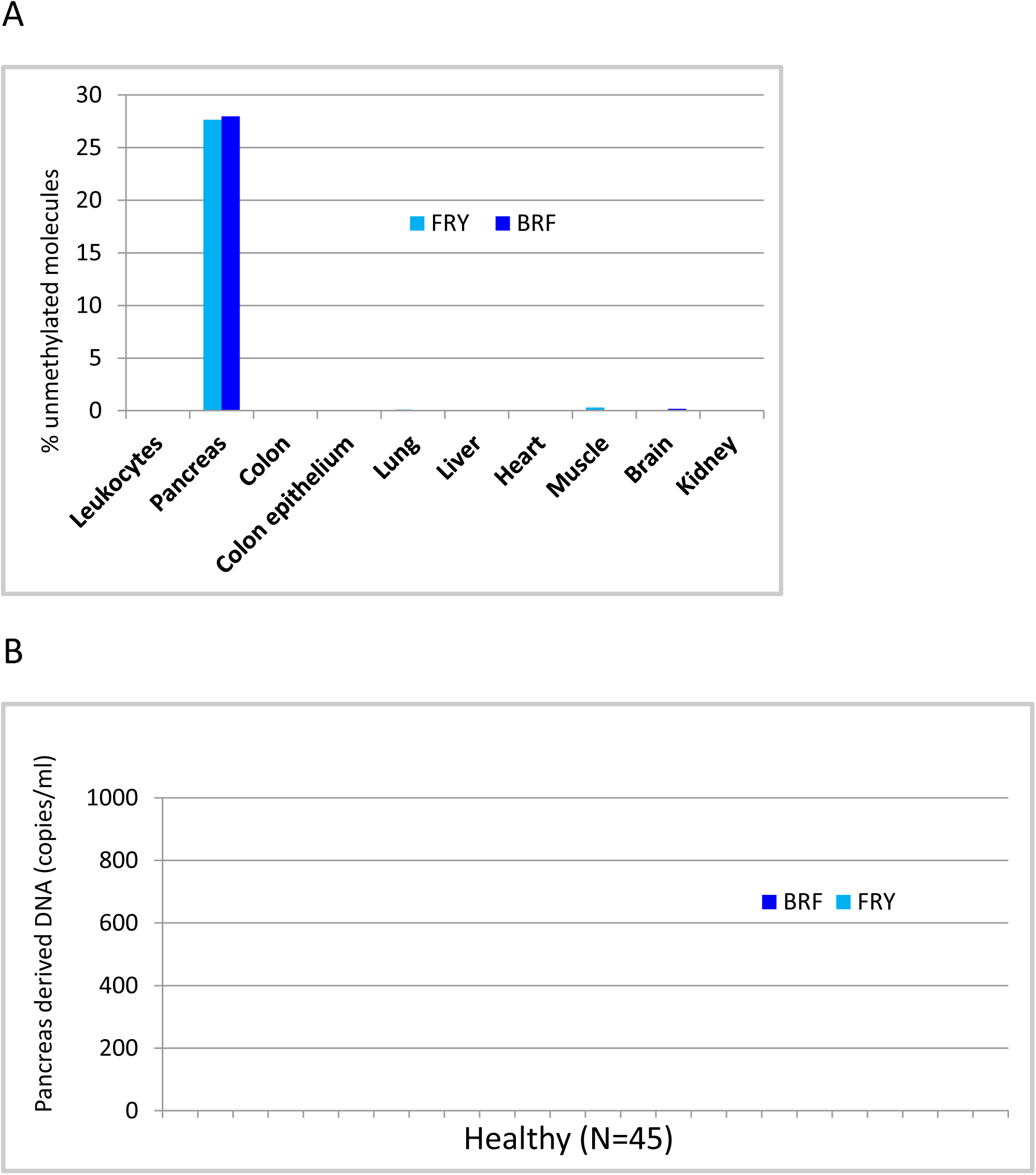
Exocrine pancreas specific markers. **A.** Methylation status of loci adjacent to the FRY and BRF genes in DNA from multiple tissues. Shown is the percentage of molecules in which all CpG sites in each amplicon were unmethylated. **B.** Concentration of pancreas derived cfDNA in the plasma of 24 healthy donors. The concentration was measured by multiplying the fraction of colon cfDNA by the concentration of total cfDNA.

